# LUstiGE, Light responsive Ustilago maydis Gene Expression: Optogenetic control of morphogenesis and pathogenesis in the corn fungal pathogen *Ustilago maydis*

**DOI:** 10.64898/2026.06.25.734638

**Authors:** Kun Tang, Micail David Müller, Lisa Hüsemann, Weiliang Zuo, Anna Rybecky, Nicole Heucken, Johannes Postma, Lasse van Wijlick, Gunther Doehlemann, Michael Feldbrügge, Matias D. Zurbriggen

## Abstract

The basidiomycete *Ustilago maydis* is a well-characterized model organism for studying pathogen-host interactions and of great interest for a broad spectrum of biotechnological applications. We set here to develop light inducible molecular tools to enable dynamic studies on signaling networks and fungi-host communication, and for metabolic engineering approaches. In particular, light-controlled, optogenetic switches provide quantitative, spatio-temporal control capabilities, are minimal invasive and reversible. We engineered two blue light-inducible LOV-domain-based gene expression switches, to up- (Blue-ON) and down-regulate (Blue-OFF) gene expression, and performed a functional characterization in sporidia and hyphae of *U. maydis*. Profiting from the dynamic control ranges and rapid kinetics, we implemented the optoswitches to control cell morphology by initiating the transition from a haploid sporidial cellular morphotype to filaments upon regulation of the levels of the polarity factor Rac1 and its constitutive active mutant Q61L. In addition to showing how expression level of effectors can be precisely regulated as an approach to understand fungi-plants interaction, we show in two proof-of-principle applications targeted control over *U. maydis* filamentous fungal invasion of plant tissue and the mechanisms of tumor formation. For this we placed under Blue-ON and Blue-OFF control two *U. maydis* effectors, See1 (<u>S</u>eedling <u>e</u>fficient <u>e</u>ffector 1) and TIN2 (<u>T</u>umor <u>in</u>ducing 2), and tumor formation was assayed on maize leaves. Taken together, this study established blue-light switches as effective tools to control morphogenesis and pathogenesis in *U. maydis*.

## Introduction

The eucaryotic model microorganism *U. maydis* has a small and well annotated sequenced genome, which allows for comparative genomic analyses and the straightforward study of signaling pathways and their components in relation to other pathogens. With the help of efficient transformation systems and the first synthetic biology tools being established ^1,2^, this organism becomes not only a suitable production host for valuable chemicals ^3,4^ but also it has turned out as a model organism to better understand eukaryotic biotrophic plant pathogens, infection dynamics as well as long-distance RNA transport ^5–9^.

Implementation of synthetic biology approaches has demonstrated to be useful to unravel and control complex cellular mechanisms. These experimental strategies and molecular tools enable the development of rapid, convenient, and urgently required biotechnological processes. For this purpose, engineering principles have been applied to design and implement molecular tools for the assembly and targeted regulation of complex natural and synthetic networks and switches^10–14^. In contrast to using chemical inducers, which involve the application of oftentimes toxic molecules, light - in turn - provides high spatial and temporal resolution and reversibility, jointly with minimal invasiveness. Over the years, a high number of light-inducible ON-/OFF-switches have been established in mammalian cell systems, plants, bacteria and yeast to control a broad range of cellular processes ^15–18^. However, besides molecular methods of manipulation such as a carbon source-regulated promoter P_CRG_ ^19^, there are no optoswitches applied in the basidiomycete fungus *U. maydis*. Implementation of light-inducible tools would provide key experimental advantages to: (i) study gene regulatory and metabolic networks, (ii) to improve and fine-tune this model system for the study of fundamental molecular, cellular and physiological mechanisms and fungus-plant interactions, (iii) contribute to widen the spectrum of biotechnological applications.

The blue light sensing light-oxygen-voltage (LOV) proteins are probably the most frequently employed light-responsive systems in eukaryotes ^15^. The LOV proteins comprise flavin derivatives as co-factor (mostly FMN, flavin-mono-nucleotide) with a blue light activatable absorption band around 450 nm. Light activated LOV-domains return to the basal, inactive state with a recovery kinetics that vary widely, spanning from seconds to hours and are widely engineered as optoswitches to manipulate gene expression, subcellular localization of proteins, enzyme activity, and metabolic pathways in prokaryotes, animals, fungi and plants _16–18,20–22._

In this study, we engineered and implemented light-inducible gene expression systems, specifically the Blue-ON and Blue-OFF optoswitches, into *U. maydis*, quantitative characterized their performance, and applied them to control cell morphology by regulating Rac1^Q61L^ gene expression. This mutant of the small GTPase, which is constitutively active, remains in a GTP-bound state, driving aberrant morphology by disrupting the regulation of the actin cytoskeleton and causing a loss of strictly polarized tip growth^23^. These tools facilitate precise spatial-temporal gene regulation in *U. maydis,* thereby enabling future studies of light controlling dynamic cellular processes. In addition, we employed the optogenetic systems to manipulate effector gene expression (e.g., See1, TIN2)^7,24^ in the *U. maydis*-*Zea mays* pathosystem, enabling light-regulated, real-time control of gene function during biotrophic infection. This approach might contribute to deepen our understanding of infection mechanisms and optimize advanced methods for bio-based applications^25^.

## Results

### Design of inducible systems for *Ustilago maydis*

*U. maydis* is an exceptionally well characterized model system for both basic and applied research. The goal of this study is to engineer, functionally characterize and implement light-inducible optogenetic systems to (i) understand fungal cell biology such as polarized growth and hyphal development, (ii) provide insight into effector proteins and host reprogramming. To address this, light-regulated gene expression systems were developed and tested in *U. maydis* cells in order to provide a toolkit for unlimited optogenetics applications (Fig. 1a). To develop a light-inducible system in *U. maydis*, synthetic elements such as inducible- promoters, reporters, and transcriptional factors are essential. A key requirement is a promoter containing target DNA sequences and the minimal cytomegalovirus promoter (P_CMVmin_), which must be implemented and tested for low basal expression in the target organism. Accordingly, the corresponding DNA-binding domains were codon-optimized, placed under a constitutive promoter and cloned into a vector for cco1 locus insertion, as described previously (Fig. 1b) ^2^.

**Figure 1.**
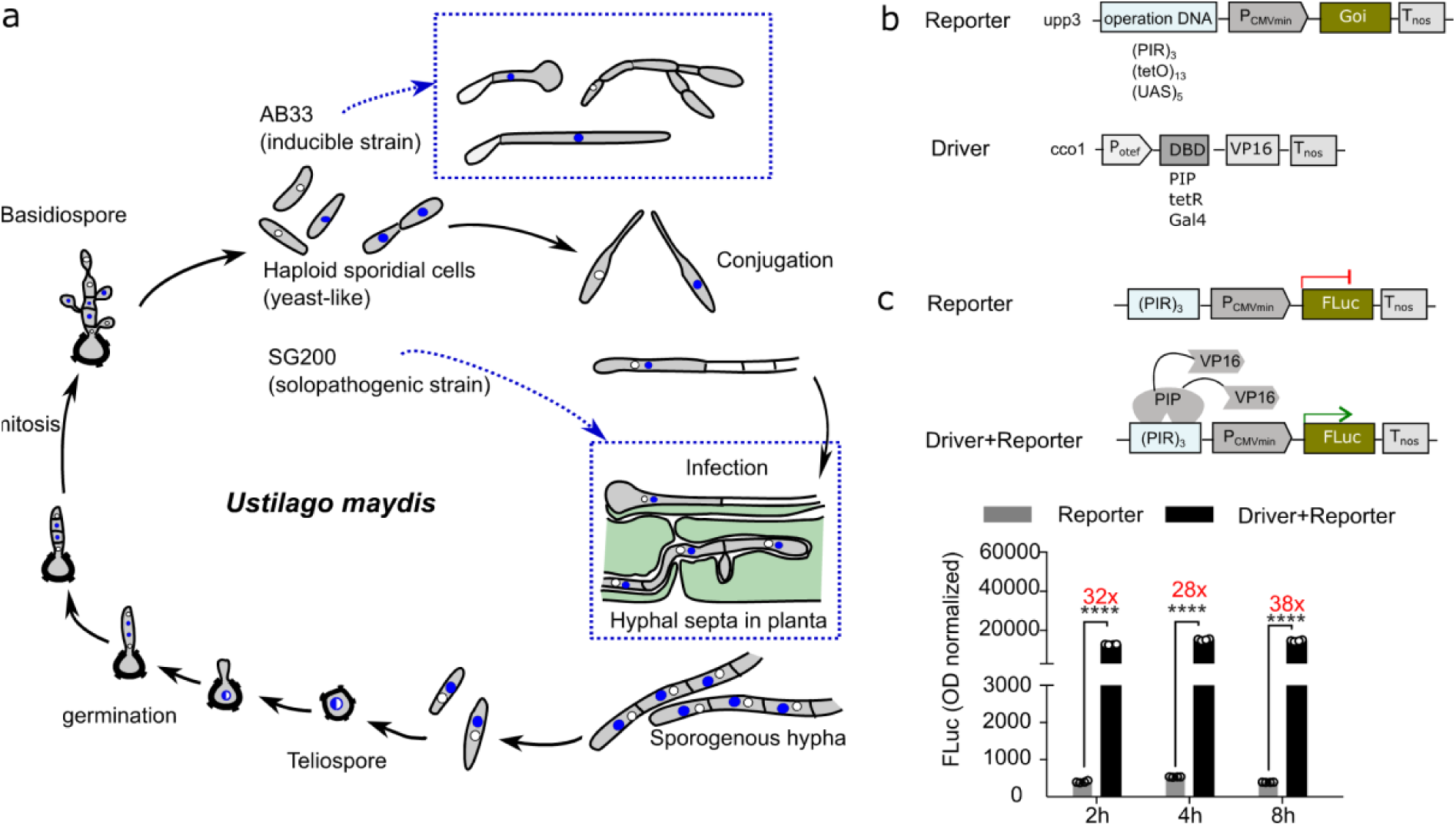
*Ustilago* cell engineering design. **(a)** Life-cycle and cell morphology of *U. maydis*. Illustration were inspired from Piepenbring^49^ and Lanvel^8^. In blue rectangles are shown the selected biological processes that are optogenetically controlled in this study: (upper) light inducible gene expression in AB33 strain for controlling cell morphology; (lower) light regulated the *Ustilago*-plant interactions with solopathgenic strain SG200. **(b)** Molecular architecture of the constructs for inducible expression of the gene of interest (Goi), and the corresponding driver plasmids for constitutive expression of the fusion protein comprising the DNA binding domain and transcriptional activator. **(c)** Schematic diagram of FLuc reporter and driver+reporter strains based on the PIP/(PIR)_3_ system, and the expression levels shown after 2 h, 4 h and 8 h. Data represent mean values with one standard deviation of four biological replicates compared with two-sided independent Student’s t-tests. ****, *p* ≤ 1e-4. Source data are provided with this paper.

While several transactivation systems exist, only the TetR-system has been shown to function in *U. maydis* ^26,27^. To further expand the molecular genetic tool box, a series of DNA-binding proteins fused to a transactivator, VP16, and orthogonal promoters consisting of the cognate DNA operator sequence and a minimal promoter were quantified for expression. The strength of these promoters was analysed by determining the activity of the firefly luciferase (FLuc) reporter, compared to a constitutive P_Otef2_ promoter (Fig. S1). The following reporter constructs were tested: (UAS)_5_ (5 repeats of upstream activating DNA sequence that efficiently binds Gal4), (PIR)_3_ (pristinamycin response element comprising 3 pristinamycin operator repeats binding PIP), and (tetO)_13_ (tetracycline response element comprising 13 tetracycline operator repeats binding TetR) ^22,28,29^(Fig. S1). We then generated stable reporter strains by homologous recombination or Cas9-mediated insertion in the *upp3* locus of the *U. maydis* genome^30^. Most of the constructs proved unsuitable due to high background expression in the absence of the transcriptional activators. The (PIR)_3_ promoter was selected as a promising candidate, exhibiting low basal signal and robust gene expression upon addition of the constitutive transcriptional activator (PIP-VP16) (Fig. 1c). In addition to the temporal control, some of these systems also allow regulation by using chemical triggers to fine-tune expression in a dose-dependent manner. When the small cognate molecular ligand is present, it should temporarily prevent the transactivator from binding to its operator resulting in a reduced expression (OFF-system^31^). To assess this potential, we measured the reporter activity under three pristinamyicin concentrations, low dose (20 µg/ml), high dose (200 µg/ml) and very high dose (2,000 µg/ml). However, the very high dose proved too toxic for *U. maydis* cell culture. Opposite to what was expected, addition of pristinamycin increased FLuc luminescence (Fig. S2)^32^. This unexpected chemical toxicity and paradoxical induction highlighted the severe limitations of using small-molecule triggers in *U. maydis*, thus prompting us to develop a more precise, non-chemical approach using light-inducible optogenetic switches.

### Blue light induced gene expression in *U. maydis* culture (Blue-ON system)

After having established an orhtogonally regulatable synthetic promoter system (PIP-VP16/(PIR)_3_, *U. maydis* strains were genomically equipped with the possibility for optogenetic gene expression control. For this, we have adapted and customized light-inducible gene expression systems that we had previously developed and tested in mammalian cells ^22,33–35^. We first engineered a blue light inducible system based on TULIPs ^21,22^. The PIP-based TULIP Blue-ON is a so-called two component system or split transcription factor, utilizing the heterodimerization of an engineered LOV2 domain from phototropin-1 of *Avena sativa* and an ePDZb domain in response to blue light (λ_max_ = 455 nm) (Fig. 2a). After generation and characterization of the strains, functionality tests of the introduced light switches were performed (Fig. 2b). The induction by blue light was determined in sporidial cell cultures with an AB33 genomic background, performing the usual quantitative characterization on the effects of light intensity, kinetics of gene expression, and the delivery of the additional co-factor FMN, in case it becomes limiting within the fungal cell (Fig. 2c) The light-dose illumination tests showed a 7-fold induction rate under 20 µmol m^-2^ s^-1^ blue light for 4 h. After 8 h illumination, a 5- fold induction was achieved (Fig. 2c). Note that addition of the flavin co-factor to the medium further amplified gene expression indicating that endogenous flavin levels might be a limiting factor.

**Figure 2.**
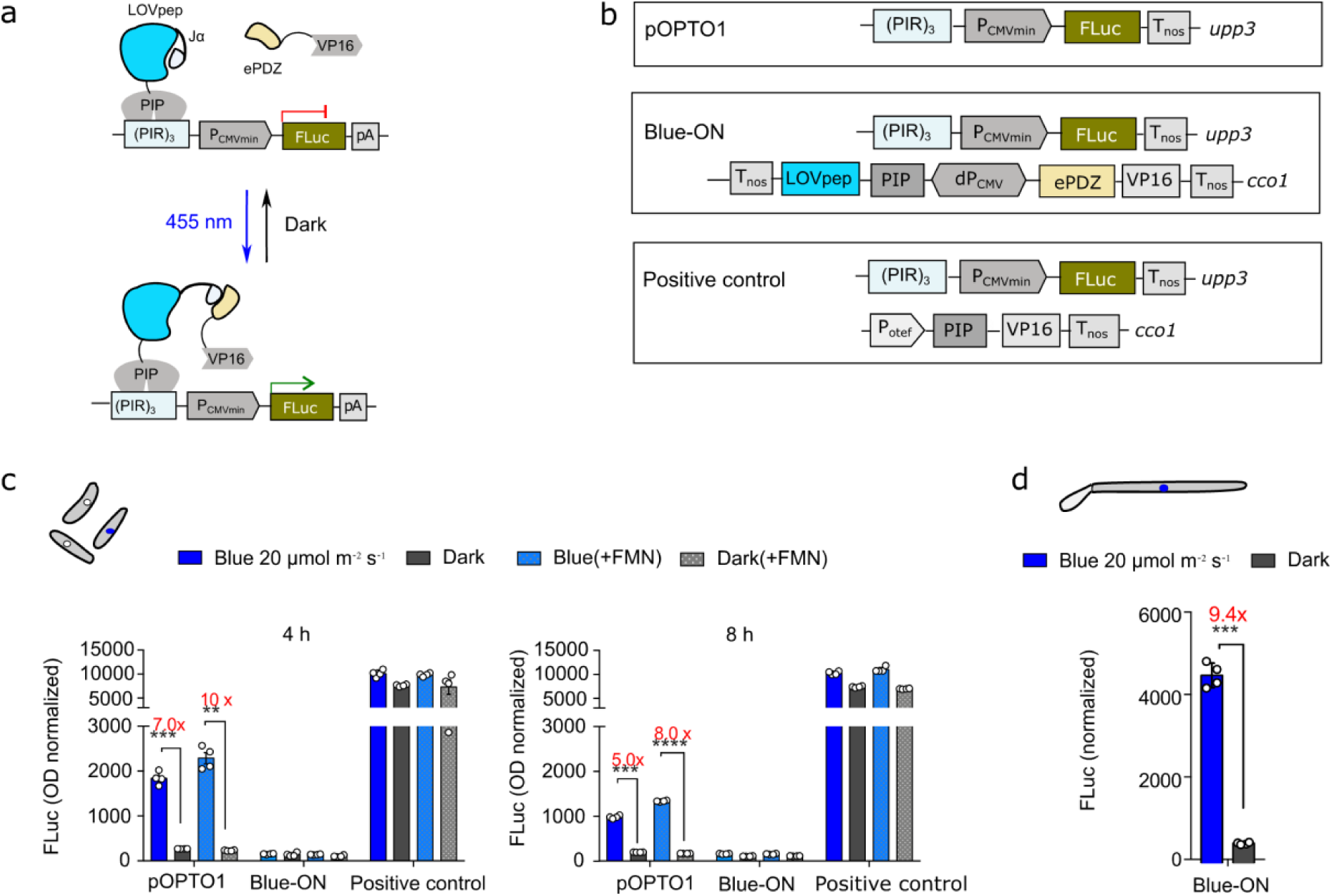
Functional characterization of the Blue-ON system in *U. maydis* cultures. **(a)** Molecular mechanism of function of the LOV-domain based Blue-ON optogenetic switch for induction of gene expression in *U. maydis*. **(b)** *U. maydis* strains engineered for the Blue-ON system. Blue-ON contains a pOPTO1 construct comprising the PIP operator sequence (PIR)_3_ upstream of a P_CMV_ minimal promoter, controlling the expression of firefly luciferase (FLuc) as a reporter gene, stably integrated in the *upp3* locus. In the *cco1* locus, a dP_CMV_ bidirectional constitutive promoter drives the expression of a fusion construct of ePDZ-VP16-NLS in the upstream direction, and a fusion of PIP-AsLOVpep in the downstream direction. The strain containing only pOPTO1 served as a negative control. A positive control for the system, containing pOPTO1 in the *upp3* locus, and PIP-VP16-NLS under the control of the constitutive P_O2tef_ promoter integrated into the *cco1* locus. **(c)** Firefly reporter activities in cultures after 4 h and 8 h in the dark or after illumination with blue light at 20 μmol m^-^² s^-1^ (λ = 455 nm). In addition, the LOV-domain co-factor FMN was added at a final concentration of 5 μM. The reporter activity in all strains was determined in cultures normalized to an OD600 of 0.5, using the firefly-based fast screening method as described in materials and methods. Data shown are means of four biological replicates. **(d)** Quantification of FLuc expression conducted in induced AB33 hyphae after 16 h. Bioluminescence was assessed in both cell-free culture supernatants and clarified whole-cell lysates after filamentation induction in nitrate minimal medium. The total reporter activity was determined through volumetric scaling and normalized to the total intracellular protein content, which was quantified using the BCA assay. Data in **c,d** represent mean values with one standard deviation of four biological replicates compared with two-sided independent Student’s t-tests. **, 1e-3<*p* ≤ 1e-2; ***, 1e-4 <*p*≤ 1e-3; ****, *p* ≤ 1e-4. Source data are provided with this paper.

To cover the complete life cycle of *U. maydis*, we measured the expression of the firefly luciferase gene in both extracellular and intracellular fractions during the dimorphic transition and within the fungal filaments, which was subsequently normalized to the absolute cellular protein mass. In cultures subjected to a photon flux density of 20 µmol m^-2^ s^-1^ of monochromatic blue light (455 nm) for 16 h following the initiation of hyphal formation by transitioning to nitrate minimal (NM) medium, a 9.4-fold induction was observed (Fig. 2d).

### Blue-light induced repression of expression in *U. maydis* culture (Blue-OFF system)

EL222 from *Erythrobacter litoralis* functions as a single molecular component in which blue light (λ_max_ = 455 nm) directly modulates the DNA-binding affinity to a sequence termed C120 (Fig. 3a). Upon illumination, both blue light switches trigger the release of a short domain, which is connected to the LOV domain through a J-alpha-helix. The EL222 protein, exposes under blue light an otherwise dark-bound HTH (helix-turn-helix) motif. This permits homo-dimerization with another EL222 protein, and subsequent binding to the target the DNA sequence (C120) ^36^.

**Figure 3.**
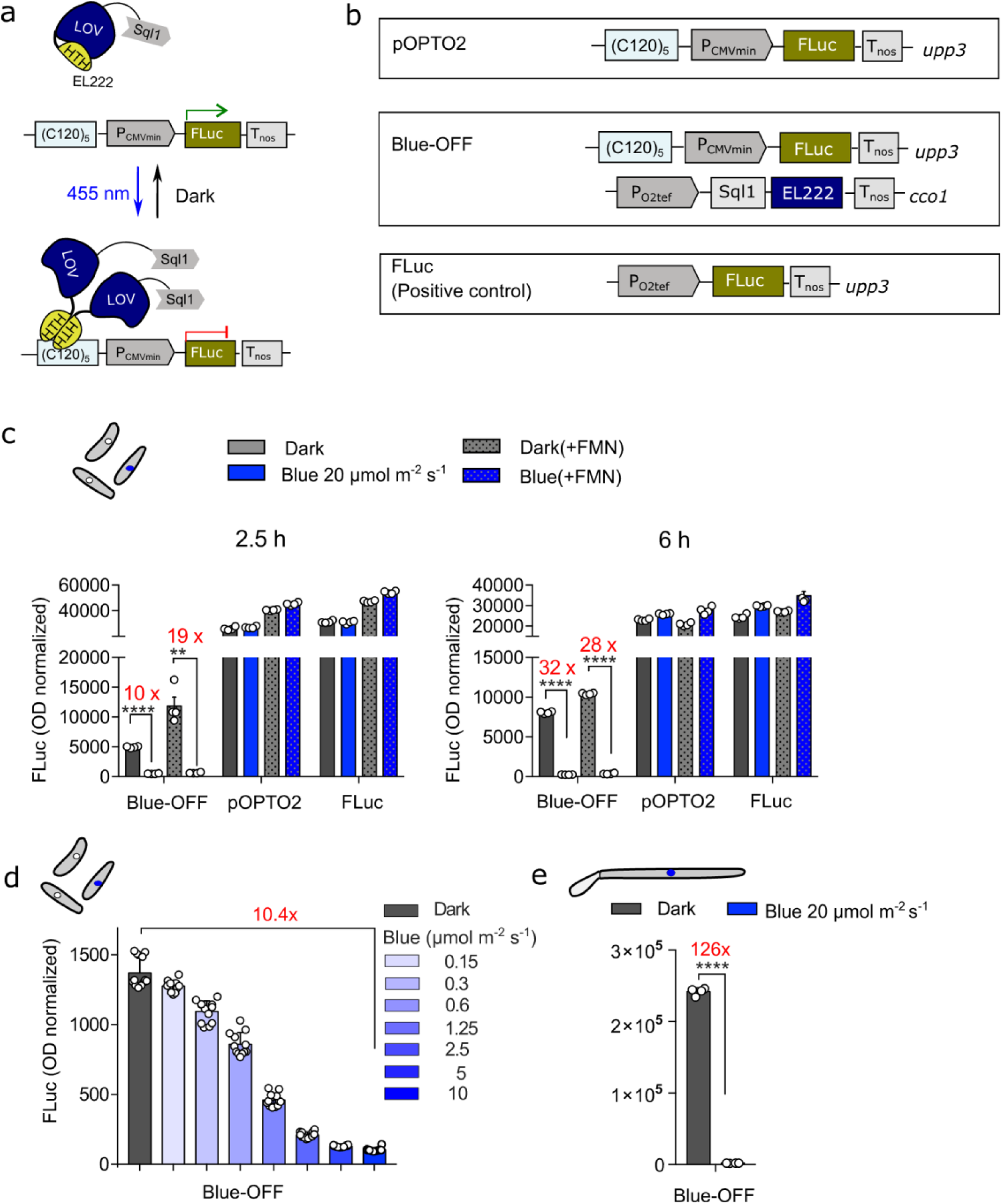
Functional characterization of the Blue-OFF system in *U. maydis* cultures. **(a)** Molecular mechanism of function of the EL222 based blue light-controlled repression of gene expression system (Blue-OFF), in *U. maydi*s. **(b)** Strain Blue-OFF was generated with two components stably integrated: i) in the *upp3* locus, 5 repeats of the EL222 operator sequence, (C120)_5_, upstream of a P_CMV_ minimal promoter, driving the expression of firefly luciferase reporter (FLuc) (pOPTO2); and ii) in the *cco1* locus, the blue light photoreceptor EL222 is fused to an *U. maydis*-derived transrepressor domain, Sql1, and placed under the control of a constitutive P_O2tef_ promoter. A control strain is with only the reporter construct pOPTO2 being integrated to the *upp3* locus. Fluc driven under constitutive promoter Potef was also tested as control. **(c)** Determination of reporter activity of Blue-OFF cultures and control strains were illuminated with blue light for 2.5 h or 6 h at 20 μmol m^-^² s^-1^ light (λ=450 nm) or kept in the dark. The LOV-domain co-factor FMN was added at a concentration of 5 μM. The reporter activity in all strains was determined in cultures normalized to an OD600 of 0.5. Data shown are means of three biological and three technical replicates. **(d)** a gradient blue light intensities (0-10 μmol m^-^² s^-1^) were tested with custom-built Light Plate Apparatus (LPA). **(e)** FLuc expression levels were quantified in AB33 hyphae following a 16-h induction period. Bioluminescence determinations were performed in both the cell-free culture supernatants and the clarified whole-cell lysates after inducing filamentation in nitrate minimal medium. The overall reporter activity was calculated by adjusting for volume and was normalized against the total intracellular protein content, which was measured using the BCA assay. Data in **d,e** represent mean values with one standard deviation of four biological replicates compared with two-sided independent Student’s *t*-tests. **, 1e-3<*p*≤ 1e-2; ***, 1e-4 <*p*≤ 1e-3; ****, *p* ≤ 1e- 4. Source data are provided with this paper.

We tested the EL222-based system exclusively with a (C120)_5_-PhCMVmin promoter with two variants of EL222, one fused to a VP16 transactivator and another to a Sql1 repressor (Fig. S3). The former leading to a Blue-ON system and the latter to a Blue-OFF. The minimal promoter alone exhibited relatively high FLuc expression (Fig. S1). In order to construct a Blue-OFF system ^34^, we replaced the transactivator with a transrepressor domain, the *U. maydis*-inherent repressor Sql1 (Fig. 3a,b amino acids 1-359). Fig. 3c shows substantial intrinsic activity of the minimal promoter, as evidenced by the comparable levels of FLuc in the dark and under blue light illumination at both 2.5 h and 6 h. Upon illumination, the repressor strongly suppresses reporter expression either with or without addition of FMN. Notably, after 2.5 h, the FLuc signal decreases to 10%, and approximately 5% when additional FMN is present. This suppression intensifies by the 6 h time point, with the signal reduced to 3 %, regardless of FMN supplementation (Fig. 3c).

The dependence of the Blue-OFF optoswitch on light intensity for effective transcriptional repression was assessed by subjecting cultures to a gradual increase in monochromatic blue light. We used the previously developed custom-built Light Plate Apparatus (LPA, materials and methods)^33,37^ to test the Blue-OFF system with light intensity range from 0 to 10 µmol/m²s. Following a 2.5-h treatment, a 10.4-fold reduction in Fluc bioluminescence was observed in sporidia (Fig. 3d).

This effect was more pronounced when evaluated in hyphae and normalized to the total protein content. Specifically, FLuc expression was approximately 126 times higher when cultures initiated for filamentation were maintained for 16 h under strict dark conditions as opposed to those exposed to blue light (Fig. 3e).

### Cell morphogenesis switch by blue light

Next, we tested whether our optogenetic tools could regulate cellular morphogenesis, focusing on the dimorphic transition from sporidia to hyphae (Fig. 4). Drawing on previous work by Mahlert et al.^23^, we engineered a Blue-OFF strain to control the expression of the constitutively active polarity factor mutant Rac1^Q61L^. Because this mutant drives isotropic cell wall expansion and the formation of large polar vacuoles, it provides a clear, distinct morphological readout for switch activity (Fig. 4a).

**Figure 4.**
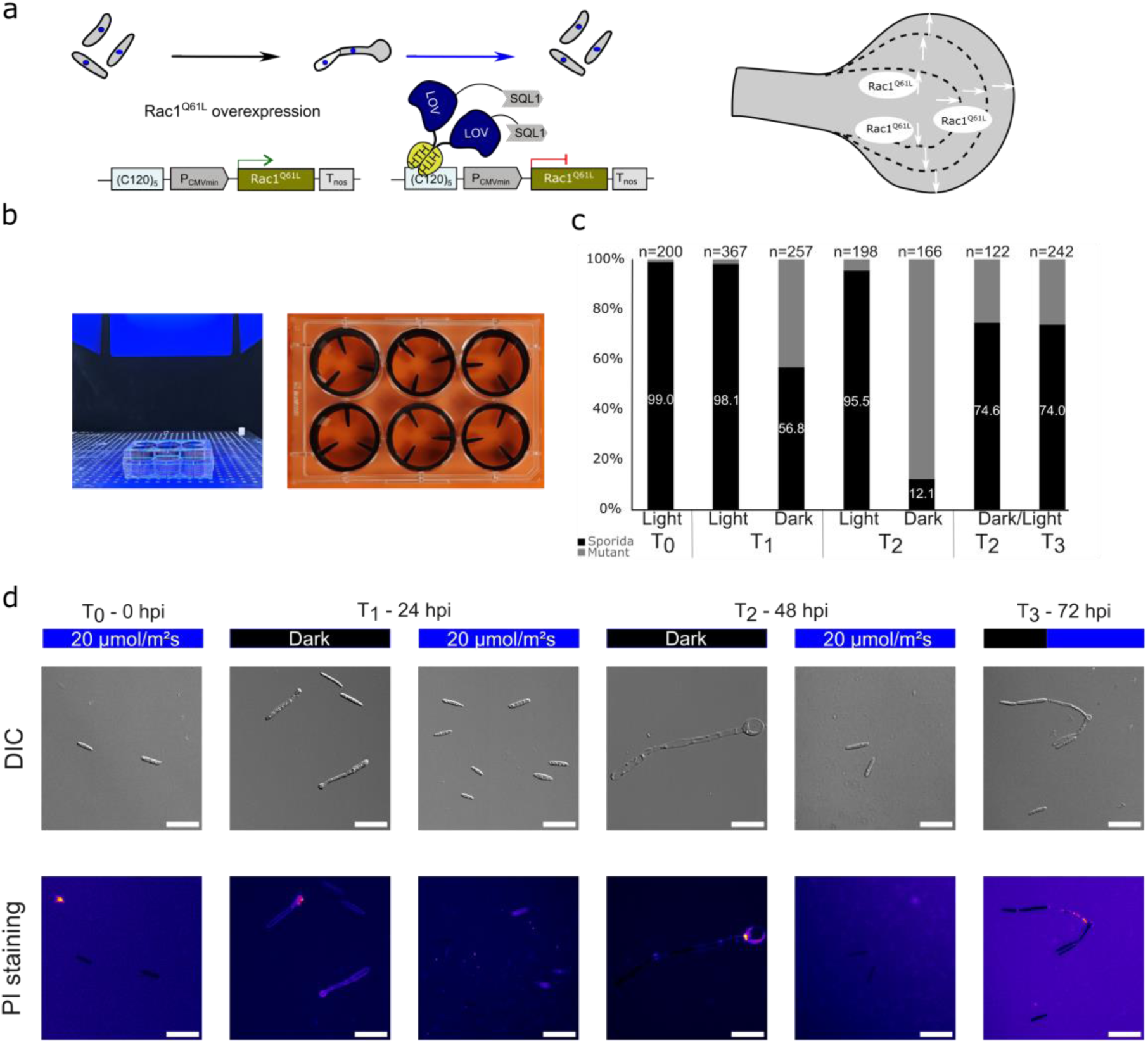
Optogenetic control of Rac1^Q61L^ expression in *U.maydis*. **(a)** Schematic representation of the Blue-OFF optoswitch controlling Rac1^Q61L^ expression. Under dark conditions the (C120)_5_ promoter remains constitutively active, driving expression of the Rac1^Q61L^ mutant and inducing the aberrant phenotype. Upon exposure to blue light (20 µmol m^-^² s^-1^), the light-responsive EL222 domain binds the synthetic promoter, thereby recruiting the endogenous *Ustilago* transrepressor domain (Sql1) to repress transcription. The Rac1^Q61L^ mutation causes isotropic cell wall expansion and the formation of large vacuoles at the poles (scheme adapted from Mahlert et al., 2006)^23^. **(b)** Cultivation platform developed for optogenetic experiments. Cultures are incubated under monochromatic blue light (λ = 455 nm) with a photon flux density of 20 µmol m^-^² s^-1^ in six-well plates containing custom 3D-printed baffled inserts. **(c)** Quantification of morphological phenotypes in dark- and light-treated cultures across specified time points. To demonstrate population reversibility, an aliquot of the dark-treated sample was shifted to blue light post-T_1_ (24 h) and assessed after an additional 24 h (T_2’_) and 48 h (T_3_) of illumination. **(d)** Representative images of predominant phenotypes in dark- and light-treated cultures. Displayed are differential interference contrast (DIC) microscopic images (top) and their fluorescent counterparts following propidium iodide (PI) staining (bottom) to visualize compromised membrane integrity. Scale bar = 20 µm.

To ensure uniform light exposure and adequate aeration during liquid culture, we grew the cells in standard six-well plates fitted with custom 3D-printed baffled inserts (Fig. 4b). This setup kept the cultures well-agitated under continuous, monochromatic blue light (λ = 455 nm) at an intensity of 20 µmol m^-^² s^-1^.

Using this platform, we assessed the morphological divergence of the engineered population under different light conditions. At the initial time point (T_0_), the cultures consisted almost entirely of normal sporidia (99.0%). When cultures were incubated in the dark to allow expression of the mutant allele, they rapidly developed the expected aberrant phenotype. After 24 h of dark incubation (T_1_), 43.2% of the cells exhibited mutant morphology. This trend continued, resulting in 87.9% heavily vacuolized cells after 48 h (T_2_). Conversely, continuous blue light illumination effectively repressed the mutant allele. Under these conditions, the cells remained predominantly sporidial (98.1% at 24 h), with only a negligible fraction showing any deformities by 48 h (Fig. 4c). As negative controls, we evaluated the wild-type strain alongside the genetic background strain used for the Blue-OFF system (which harbors the Δcco1 and Δupp3 deletions). As expected, these controls remained exclusively as sporidia under all light conditions and time points (Fig. S4).

To determine whether this extreme isotropic expansion affected cell viability, we performed propidium iodide (PI) staining to assess cell membrane integrity and identify compromised cells. The PI staining provided clear evidence that the cell membranes of the enlarged Rac1^Q61L^ mutants were severely compromised, ultimately resulting in cell lysis and death (Fig. 4d).

Despite the lethality of the mutant phenotype at the individual cell level, we tested the control capabilities of the Blue-OFF switch to see if the culture could recover. We shifted an aliquot of the dark-treated, morphologically aberrant cultures (post-T_1_) back under blue light illumination. Strikingly, we observed a clear population-level shift back towards sporidial dominance, reaching approximately 74% normal sporidia after both 24 (T_2_) and 48 h (T_3_) of re-illumination (Fig. 4c). Microscopy of these recovering cultures revealed the presence of several elongated, dying mutant cells from which new, healthy sporidia were actively emerging through budding (Fig. 4d). This demonstrates the rapid and precise control capabilities of the Blue-OFF system in regulating *U. maydis* cellular processes, even following the induction of a terminal phenotype.

### Optogenetic control of the *Ustilago* Seedling Efficient effector 1 for inducible tumor formation

Achieving tight, precise control over tumor formation is a long-sought goal in both developmental biology and pathology, as natural tumorigenesis is often stochastic, irreversible, and influenced by complex, poorly resolved host-microbe or cell-autonomous signals ^8,38–40^. Inducible systems that allow spatiotemporally restricted tumor initiation would enable researchers to dissect early transformation events, test virulence windows, and model tumor formation with unprecedented resolution. For instance, the control of effector deployment with light-induced expression or secretion, might facilitate the study of timing and dosage effects on host responses. We set out to implement optogenetic control over the leaf-specific effector See1(<u>S</u>eedling <u>e</u>fficient <u>e</u>ffector 1)^41,42^. See1 activates host DNA synthesis to directly promote tumor formation in bundle sheath cells (Fig.5a).

To achieve light-inducible regulation of the *U. maydis* core effector See1, we implemented two optogenetic systems: the LOVpep/ePDZ based Blue-ON switch (Fig. S5a) and the EL222-based Blue-OFF switch (Fig. 5b). The coding sequence of See1 was fused to FLuc to enable non-invasive real-time quantification of expression. For the Blue-ON system, See1-FLuc was placed under the control of the (PIR)_3_ promoter, generating the reporter strain (s1836) and Blue-ON-See1 strain (s1877) (Fig. S5a,b). A positive control strain, designated s333B, with PIP-VP16 inserted into the *cco1* locus was also generated (Fig. S5a). For the Blue-OFF approach, we used the EL222-Sql1 system: strain s1874 was constructed by integrating Potef-EL222-Sql1 into the *cco1* locus and (C120)_5_-P_CMVmin_-See1-FLuc into the *upp3* locus. A non-light-responsive control strain (s1835) carrying (C120)_5_-P_CMVmin_-See1-FLuc alone in *upp3* was also generated (Fig. 5c). All strains were derived from the solopathogenic progenitor SG200.

**Figure 5.**
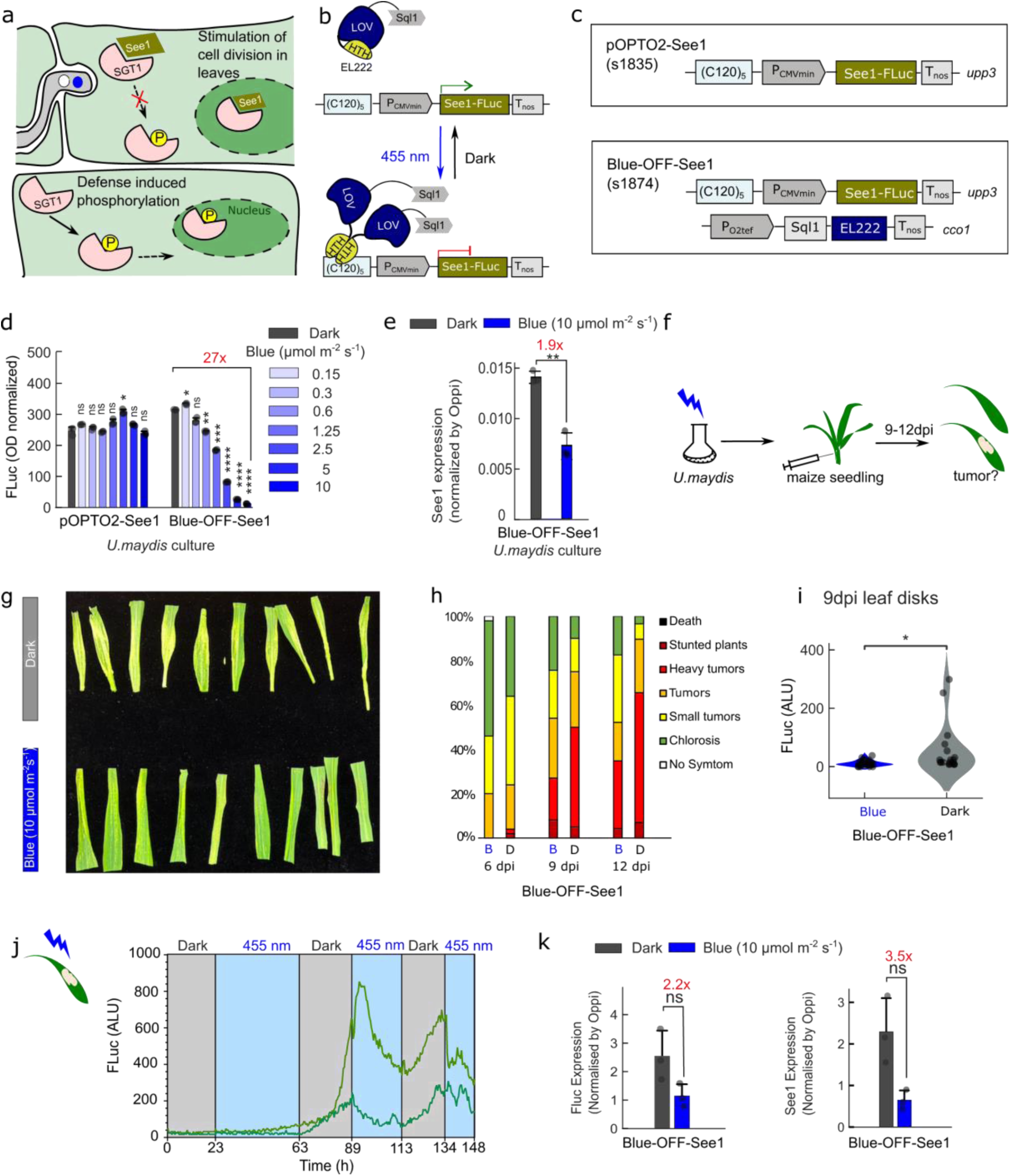
Optogenetic control of tumor formation in maize leaves using Blue-OFF regulation of See1 expression in *U. maydis*. **(a)** Schematic of tumor formation in maize seedlings. See1 interacts with defense-induced phosphorylated maize SGT1 (suppressor of G2 allele of *Skp1*) to prevent activation of downstream immune responses (upper cell). In the absence of See1, SGT1 phosphorylation occurs, leading to immune activation (lower cell). **(b)** Functional molecular description of the Blue-OFF mediated optogenetic regulation of See1 gene repression, and **(c)** strains engineered. **(d)** FLuc activity (normalized to OD600) determined at OD600 = 0.8. Blue light intensity gradients (0 - 10 µmol m^-^² s^-1^) were tested for the Blue-OFF-See1 (s1874) and control (s1835) strains. **(e)** RT-qPCR quantification of See1 mRNAs in exponentially growing *U. maydis* strains. **(f)** workflow of See1 infection maize seedling. **(g)** Representative leaf photographs showing phenotypic differences betweenSG200- and Blue-See1-infected GB maize cultivars at 12 d post-infection (dpi). **(h)** Virulence phenotypes of the plants injected with dark(D) or blue-light-treated(B) *U. maydis* Blue-OFF- See1 strains, samples were evaluated at 6 dpi, 9 dpi and 12 dpi. **(i)** FLuc activity from fresh leaf disks collected at 9 dpi. (j) FLuc dynamics (time-response) in 9-dpi leaf disks over 148 h under three cycles of dark/blue light; illumination conditions are indicated. **(k)** FLuc and See1 expression quantified by RT-qPCR in leaves infected with Blue-OFF-See1. Data in **e, f** and **k** represent mean ± SD from three biological replicates. Data in **i** represent 3 leaf disks from one infected leaf and 4 infected leaves were collected. Significance was determined using a *t*-test for the data in **e, i, k** and ANOVA test for data in **f**.; ns, not significant; *, 1e-2<*p* ≤ 0.05; **, 1e-3<*p* ≤ 1e-2; ***, 1e-4 <*p* ≤ 1e-3; ****, *p* ≤ 1e-4. Source data are provided with this paper.

We first tested expression in the *U. maydis* strains under different light conditions, and found that white light (117 µmol m^-^² s^-1^) and monochromatic blue light (455 nm, 10 µmol m^-^² s^-1^) induced equivalent levels of FLuc expression (Fig.S5c,d). Using the Blue-OFF strain, a blue light intensity gradient (0-10 µmol m^-^² s^-1^) showed that 5 µmol m^-^² s^-1^ sufficed to suppress most FLuc expression, whereas 10 µmol m^-^² s^-1^ resulted in tighter repression with a dynamic ratio of 27-fold (Fig.5d). Light-regulated See1 expression was further quantified by RT-qPCR in axenic culture (Fig.5e).

Subsequently, we evaluated the optogenetic regulation of plant infection. Light-treated axenic cultures were injected into leaves of 7- to 9- day-old seedlings as described previously^2,7,41^. The inoculation site was positioned in the middle of the plant’s stem, surrounded by three leaves, which partially limited light exposure to the pathogen. Leaves were collected from 3 to 15 dpi (Fig. 5f). Successful leaf infection was observed from both the Blue-ON and -OFF systems. However, there was no noticeable severe tumor formation observed from the Blue-ON system (Fig. S5c). This concurs with previous reports, in which ectopic overexpression of the effector gene See1 did not increase the virulence of *U. maydis* in maize leaves ^41^. In contrast, the Blue-OFF system showed functional light response in the infected plant leaves (Fig. 5g). Although plants injected with blue-light-treated and dark cultures were both grown under long-day photoperiods, tumor scoring from 6 to 12 dpi revealed a higher tumor formation rate in leaves inoculated with *U. maydis* cultures incubated in the dark (Fig. 5h). Thus, we assume the higher expression level from axenic cultures would be enough to trigger tumor formation.

Building on our previous work using Fluc expression in *U. maydis* infected maize leaves ^2^, here we attempted optogenetic control and quantification of FLuc expression directly from the infected leaves. Leaves infected with the Blue-OFF-See1 strain were collected at 9 dpi and the luminescence was determined (Fig.5i). The leaf disks were further illuminated over a total time period of 148 h under alternating blue light/dark cycles and luminescence were measured every 30 min (Fig. 5j). FLuc expression was elevated in the dark (time period 63 h up to 89 h). The following blue light period (89 h up to 113 h) leads to a reduction of FLuc. This pattern is repeated with slightly lower intensity during the following light-dark periods. Finally, qPCR analysis of 9 dpi leaves confirmed light-dependent expression of both FLuc and See1.

### Optogenetic induction of *Ustilago* effector TIN2 for anthocyanin biosynthesis

In addition to control and monitor *Ustilago* effector activity, we set up to also study the host plant response. For this, we selected TIN2 as a target for optogenetic control, as its expression leads to a readily visible anthocyanin readout in maize tissue. TIN2 is a secreted effector of *U. maydis* which stabilizes a maize kinase that promotes anthocyanin biosynthesis while reducing lignin biosynthesis during infection ^43,44^. Of particular interest is that the deletion of TIN2 results in a reduced size of *Ustilago-*induced tumors, and, most obviously, a complete loss of anthocyanin production in the infected maize tissue, which is a typical feature in wildtype infections ^45^ (Fig.6a).

We first engineered the TIN2 knock-out strain s798, which induces a consistent phenotype with no anthocyanin being produced in the leaf upon infection ^46^. To explore optogenetic control of TIN2-mediated tumor phenotypes, we then engineered the Blue-ON construct and introduced it into the background strain s798. The resulting strain s2575 demonstrated that the implementation of the light switches alone (no TIN2 reporter inserted) in *U. maydis* does not affect the biotrophy, neither interact with the plant anthocyanin biosynthesis.

We next generated strain s2583, in which the Blue-ON switch was integrated into the *cco1* locus and opto-TIN2 into the *upp3* locus (Fig.6b). Light-dependent TIN2 expression was confirmed in axenic *U. maydis* cultures illuminated with blue light (Fig.6c) and infection progression was monitored over a two weeks period (Fig.6d). The extraction and quantification of anthocyanin revealed a distinct red pigmentation in the group of leaves infected with the blue-light treated Blue-ON-TIN2 *U.maydis* culture (Fig.6e). In the anthocyanin biosynthetic pathway (Fig. 6f), expression of late biosynthetic genes determines the quantitative variation in anthocyanins. Transcript levels of late biosynthetic genes decrease during later stages of ripening when discoloration occurs^47^. Specifically, F3H and DFR exhibited the highest fold increases in the plants infected with blue-light illuminated *U. maydis* culture ^48^ (Fig. 6g,h).

**Figure 6.**
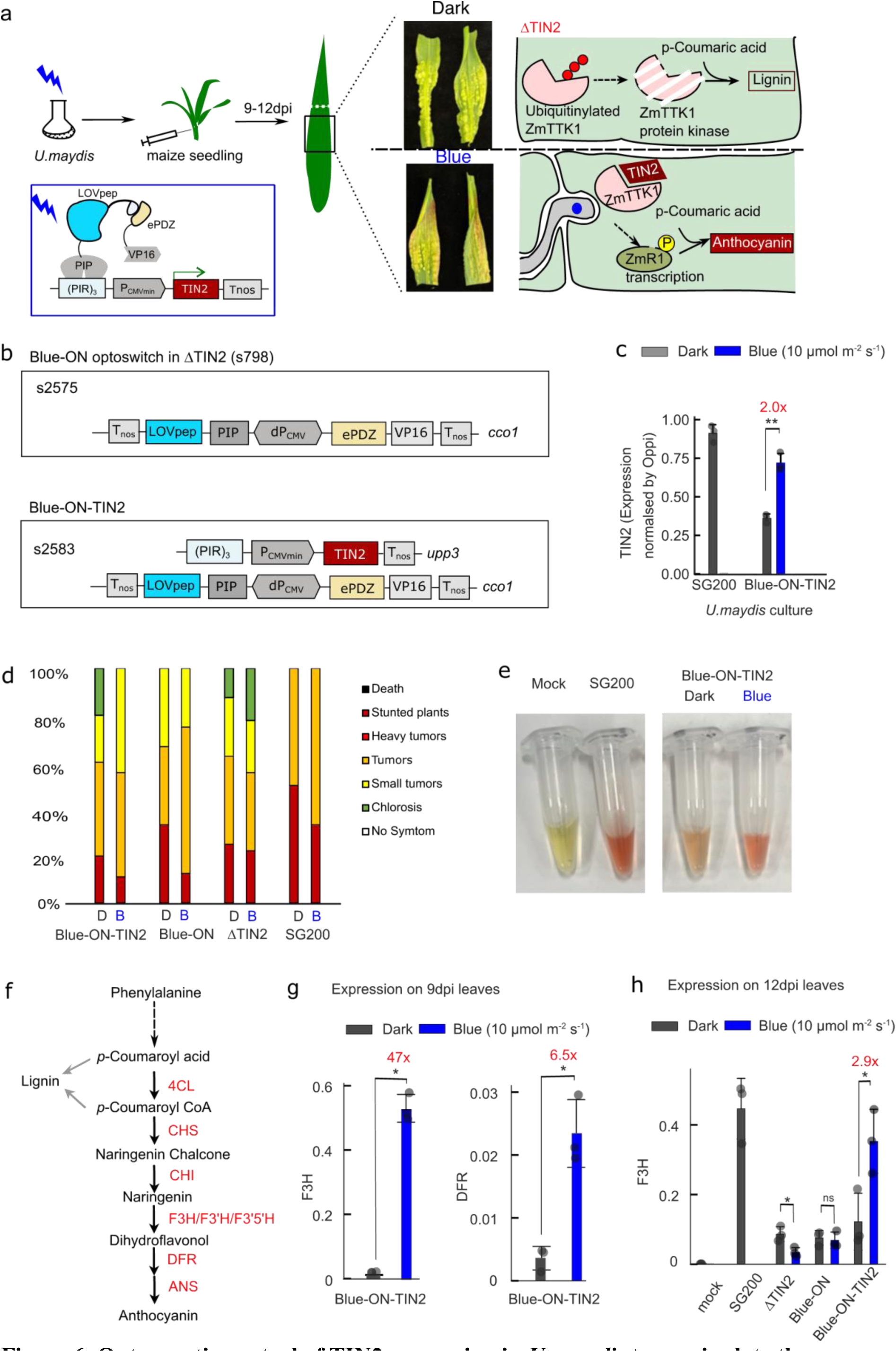
Optogenetic control of TIN2 expression in *U. maydis* to manipulate the anthocyanin biosynthesis in plants. (a) Schematic working mode of *U. maydis* expressed TIN2 in maize seedlings upon infection upon optogenetic expression control. The deletion of TIN2 results in a complete loss of anthocyanin production in the infected maize tissue (upper cell). TIN2 stabilizes a maize kinase that promotes anthocyanin biosynthesis while reducing lignin biosynthesis during infection (lower cell). The representative images of the 12 dpi-leaf inoculated with light- or dark-treated Blue-ON-TIN2 *U. maydis* culture. Tin2 was induced by blue light illumination in *U. maydis* culture, which upon inoculation induced the anthocyanin biosynthesis in the plant tissue. This leads to a red coloration observed in the leaves. **(b)** Molecular architecture of the constructs used and **(c)** Host strains description equipped with Blue-ON controlled TIN2 expression in *U. maydis*. **(d)** Tumor scoring of Blue-ON-TIN2 infected (s2583) leaves at 12 dpi. **(e)** Anthocyanin extraction and absorption spectrum from 12 dpi leaves. **(f)** Schematic model of the flavonoid biosynthetic pathway in maize. The enzymes involved in the discrete biosynthetic steps are shown in red. CHS, chalcone synthase; CHI, chalcone isomerase; F3H, flavanone 3-hydroxylase; DFR, dihydroflavonol 4-reductase; ANS, anthocyanin synthase. Anthocyanin biosynthesis and selected genes (F3H and DFR) were quantified in 9dpi (g) and 12 dpi (h) between leaves injected with dark and blue-light treated *U. maydis* culture. Values represent the mean of three biological replicates with two technical duplicates each. Error bars represent the SEM. Statistical significance was determined using a t-test for the data in **c,g** and **h**.; ns, not significant; *, 1e-2<*p*≤ 0.05; **, 1e-3<*p*≤ 1e-2; ***, 1e-4 <*p* ≤ 1e-3; ****, *p* ≤ 1e-4. Source data are provided with this paper.

## Discussion

The strong infectivity of *Ustilago maydis* in maize, together with its fully sequenced genome, has established this fungus as a well-characterized model for studying pathogen-host interactions. Detailed analyses aim to understand signaling pathways, penetration mechanisms, and key infectivity effectors. However, the few existing experimental approaches available to manipulate signalling and metabolic networks suffer from limitations such as invasive chemical treatments, poor spatio-temporal control due to diffusion, and potential toxicity.

In this study, we established a suite of optogenetic tools to achieve precise, reversible, and spatiotemporally controlled induction of *U. maydis* genes and effector activity, thereby overcoming major limitations of constitutive or chemically inducible systems in fungal phytopathology. Our results demonstrate, first, the successful design of a light-inducible gene expression switch adapted to *U. maydis*, which proved functional in liquid culture under blue light illumination, i.e. TULIP-based Blue-ON^21,22^ (Fig.2,3). In addition, we also developed a Blue-OFF configuration with EL222, offering fast deactivation and reduced background activity.

Extending beyond transcriptional control, we achieved optogenetic regulation of cellular morphogenesis, specifically the dimorphic transition from sporidia to hyphae, which is critical for *U. maydis*. Building on the foundational discoveries of Mahlert et al.^23^, we engineered a strain in which the constitutively active mutation of Rac1 is regulated by the Blue-OFF system. Much of our foundational understanding of fungal cell polarity stems from the classic budding yeast model *Saccharomyces cerevisiae*, which, notably, lacks a true Rac1 ortholog entirely and relies strictly on Cdc42 as the master regulator of symmetry breaking and polarized tip growth ^50,51^. In contrast, more complex dimorphic fungi utilize both GTPases; for instance, in the closely related human fungal pathogen *Candida albicans*, Rac1 and Cdc42 play distinct, non-redundant roles during hyphal development and morphogenetic switching^52^. Similarly, *U. maydis* requires both signaling modules to orchestrate filamentous growth ^23^. Furthermore, the spatiotemporal dynamics of polarity establishment in these fungi are highly distinct from mammalian systems; rather than mutually organizing a ’front’ and ’back’ over long distances via mechanical tension, fungal Rho and Rac cooperate locally at the apex to drive unidirectional growth^53^. By utilizing the Rac1^Q61L^ mutant, which disrupts this fine-tuned local coordination and induces isotropic cell wall expansion alongside the formation of large polar vacuoles, we established a readily distinguishable morphological readout. Crucially, our data highlights the rapid and precise reversibility of the Blue-OFF system. Upon re-illuminating dark-treated, morphologically aberrant cultures with blue light, we observed a clear population shift back toward sporidial dominance. We hypothesize that while the Rac1^Q61L^mutation severely compromises cell membrane integrity, ultimately leading to cell death, as evidenced by our propidium iodide staining, the phenotype remains reversible at the population level. This likely occurs through the budding of new, non-mutant daughter cells from the surviving elongated mutant cells. Further investigation, such as tracking the fate of individual aberrant cells via microfluidics or staining the nuclei of purported new daughter cells, will be necessary to fully confirm this mechanism.

Consequently, the establishment of light-controlled filamentation in this work not only recapitulates natural infection-related differentiation but also enables the uncoupling of filamentous growth from effector secretion in future mechanistic studies. Most importantly, in the two selected proof-of-principle applications of the opto-tools we aimed at controlling effectors. Light-repressed See1 expression reduced tumor formation in maize leaves (Fig. 5), while TIN2 activation drove ectopic anthocyanin biosynthesis, a visual, non-destructive reporter of effector activity (Fig. 6). In particular, in the plant pathogen experiments with TIN2, the anthocyanin readout proved particularly valuable as it allows real-time tissue monitoring of effector function without destructive sampling. A key advantage of optogenetics is the ability to achieve spatial control of gene expression. Using precisely timed illumination regimes, our tools provide a promising approach to real-time monitor effector expression levels and to dissect their direct and indirect effects on plant defense regulation. In Fig. 5, a FLuc-See1 fusion under Blue-OFF control was expressed in infected maize leaves, demonstrating that this optogenetic switch remains dynamic, reversible, and quantifiable within living plant tissue throughout biotrophic interaction. Taken together, these findings establish a versatile optogenetic platform in *U. maydis* that enables tunable, reversible, and spatially resolved effector studies. Unlike chemical inducers (e.g., tetracycline), light avoids pleiotropic effects, tissue penetration gradients, and washout delays, i.e. reversibility. The ability to initiate tumor formation at defined times and locations opens avenues to dissect early host reprogramming events that are normally masked by progressive, asynchronous infection. In the current study, while our blue light-based optogenetic switches enabled inducible effector control, this system has notable limitations: blue light can induce phototoxicity, cause plant growth defects, and trigger unintended host stress responses. Furthermore, blue light suffers from poor tissue penetration and is difficult to apply locally due to the rapid dark reversion and also fast development of maize seedlings, which outgrows confined illumination zones. Looking forward, we aim to engineer bi-stable toggle switches such as the red-light-based optogenetic switches (e.g., PULSE, plant usable light-switch elements)^54^ for *U. maydis*, as red light offers deeper tissue penetration, lower phototoxicity, and greater compatibility with normal plant growth cycles (light/dark) and simultaneous imaging.

In this study, the successful optogenetic manipulation in *U. maydis* demonstrates the utility of light-based control for dynamic gene regulation in diverse host-microbe systems. Future work will integrate these optogenetic modules into co-culture systems or live-imaging setups to map effector delivery dynamics and host responses with single-cell resolution. Such precision will allow us to dissect effector trafficking, local host target engagement, and the spatiotemporal dynamics of tumor initiation at near-molecular resolution. Ultimately, this platform would help transforming *U. maydis* from a genetically tractable pathogen into a light-programmable model for tumor initiation, potentially informing principles of oncogenic growth control across eukaryotes.

## Methods

### Plasmid construction

All plasmids constructed in this study were generated using standard restriction and polymerase chain reaction (PCR)-based assembly cloning methods and documented using GMOCU ^56^. The *U. maydis* codon optimised gene fragments of TULIP, EL222 were synthesis by GeneArt (Thermo Fisher Scientific), as plasmid DNA which we transformed into *U. maydis* strain AB33 or SG200 to generate stable transgenics. Plasmid details are provided in Supplementary Tables S1.

### Generation of stable transgenic strains

Strains that were constructed in this work are listed and described in Table S2. We engineered AB33 strains for the quantitative reporter assay ^57^ and the Rac1^Q61L^experiments as well as SG200 strains for *U. maydis* virulence assay ^58^. *U. maydis* stable transformation was performed with linearized plasmids using homologous recombination with the *cco1* or *upp3* locus according to Bösch et al. (2016)^59^. For sequence integration in the *cco1* locus, plasmids were digested with the restriction enzyme SspI. For integration in the *upp3* locus, plasmids were digested with SwaI. Samples that were digested with SwaI were incubated at 65 °C for 20 min to inactivate the enzyme. For SspI digested samples the plasmid DNA was cleaned-up and isolated using a DNA clean up kit (NEB).

The TIN2 related strains were generated by CRISPR-Cas9 mediated knock-in as described before^60^. In brief, the sgRNAs were designed by chopchop ^61^ to target the *upp3* and *cco1* locus and were subsequently cloned into the pCas9HF1 vector by Gibson assembly ^30^. The protoplasts of *U. maydis* were co-transformed with CRISPR-Cas9 plasmid and donor plasmid (1.5 µg each). The resulting mutants were confirmed by means of southern blot analysis. sgRNA oligo sequence: sgRNA_K_cco1:

CAAAATTCCATTCTACAACGCGTTGGTACGAAAGCGCGGGTTTTAGAGCTAGAAA TAGC; and sgRNA_K_upp3: CAAAATTCCATTCTACAACGATACGGGTACAACGCTCATGTTTTAGAGCTAGAAA TAGC.

Correctness of all strains was confirmed by genotyping PCR and/or southern blot analysis ^62^.

### *U. maydis* Cultivation

Glycerol stocks of *U. maydis* strains were streaked onto Complete Media (CM) agar plates supplemented with 1% glucose and maintained at 28°C for 2 to 3 days. These served as the initial material for inoculating primary cultures in liquid CM media with 1% glucose. The cultures were incubated as batch cultures in either baffled glass flasks or six-well plates with 3D-printed baffled inserts at 28°C and 150 to 200 rpm. Hyphal induction of AB33-based strains was achieved by replacing CM with Nitrogen Minimal (NM) media. Fungal sporidia cultures in NM media were adjusted to a final OD_600_ of 0.5 to 0.8 and cultivated in six-well plates with 3D-printed baffled inserts at 28°C and 150 rpm in a incubator (KB_E6, Binder GmbH, Tuttlinen, germany) for approximately 16 hours.

### Luminescence Determination

Luminescence was determined using a Berthold Technologies Centro XS3 LB960 Microplate luminometer as previously described ^2^. For FLuc assays in sporidial cell culture, 80 μL of the culture was transferred to white 96-well assay plates and 20 μL of firefly substrate was added as described above. The luminescence was measured in a Berthold Technologies Centro XS3 LB960 Microplate luminometer. The OD600 measurement was performed on a CLARIOstar plate reader (BMG LABTECH), and the obtained values were normalised to an OD600 of 0.5.

For the Light-Plate-Apparatus (LPA^37,33^)-based intensity tests primary cultures were inoculated and incubated for approximately 24 h at 28°C in test tubes placed on rotating wheels. The experimental main culture was adjusted to an OD600 of 0.1-0.2, from which 1 mL was transferred to each well of an entire black (transparent bottom) 24-well plate. Subsequently, the LPA was mounted on an orbital shaker set to 150 rpm within an incubator (KB_E6, Binder GmbH, Tuttlinen, germany) maintained at 28°C, where the cultures were incubated at the indicated time points.

### Bioluminescence Reporter Assay and Biomass Normalization

To quantify gene expression during the dimorphic transition and in filaments, firefly luciferase (FLuc) activity was measured in both extracellular and intracellular fractions of *U. maydis* strains Blue-ON and Blue-OFF (AB33 genetic background) hyphal cultures. Filamentous growth was induced in 4 ml of nitrate minimal (NM) medium for 16 h. Cultivation was conducted in 6-well cell culture plates with ad-hoc 3D-printed baffled inserts on an orbital shaker (150 rpm) in a custom-made incubator with LED-based illumination at 28°C. Plates containing dark-treated samples were tightly wrapped in aluminium foil, whereas plates containing light-treated samples were exposed to monochromatic blue light (455 nm, 20 µmol/m²s).

#### Fraction Separation and Whole-Cell Lysis

Cultures (4 mL) were harvested by centrifugation at 10,000 rpm for 5 min at 4°C. To capture the extracellular reporter fraction (E), a 1 mL aliquot of the cell-free supernatant was collected, transferred to a sterile 1.5 ml microcentrifuge tube on ice, and the remaining supernatant was then discarded.

The intact pellet was washed in 1 ml ice-cold phosphate-buffered saline (PBS) and centrifuged at 10,000 rpm for 10 min at 4°C. Subsequently, the supernatant was completely removed, and the pellet was resuspended in 800 μL of filter-sterilized customized 1× Passive Lysis Buffer (10 mM Tris-HCl pH 8.0, 0.1% v/v Triton X-100) supplemented with a 1× cOmplete™ Mini, EDTA-free Protease-Inhibitor-Cocktail (#11836170001, Roche).Additionally, approximately 130 mg of glass beads were added to each tube for cell lysis. The samples were immediately snap-frozen in liquid nitrogen to preserve enzyme stability and structural integrity. Cellular lysis was performed via cryo-mechanical shattering using a benchtop vortex mixer at maximum speed for 5 min in sequential 1 min intervals (1 min vortex, followed by 1 min on ice). The resulting homogenate was clarified by centrifugation at 10,000 rpm for 15 min at 4°C to remove cellular debris and glass beads. Subsequently, the supernatant was transferred to fresh 1.5 ml microcentrifuge tubes, yielding a clear intracellular lysate fraction (L), which was kept on ice.

#### Luciferase Activity Measurements

Bioluminescence was quantified in 96-well white, flat-bottom assay plates using a Centro XS³ LB 960 Microplate Luminometer (Berthold Technologies). For both fractions (E and L), 80 μL technical quadruplicates (E) and quintuplicates (L) were transferred into individual wells. 20 µl of D-Luceferin Firefly substrate (# L-8200, Biosynth) was added per well and measurement (0.1 second intervals with 29 repeats) was immediately started.

#### Protein Quantification and Specific Activity Calculation

To normalize bioluminescence to total cellular biomass, the protein concentration of the intracellular lysate (L) was determined using a Pierce™ BCA Protein Assay Kit (Thermo Fisher Scientific). Quadruplicate samples (25 μL) were incubated with 200 µL of the bicinchoninic acid working reagent for 40 min at 37°C in a dry air incubator to compensate for the initial thermal lag, followed by cooling for 5 min at room temperature. Blank-corrected absorbance was measured at 562 nm using a CLARIOstar plate reader (BMG LABTECH). Absolute protein concentrations (μg μL^-1^) were calculated against a linear calibration curve generated from duplicates of bovine serum albumin (BSA) standards diluted in identical 1× Passive Lysis Buffer.

Total culture gene expression was calculated by scaling the raw RLU values to the absolute fraction volumes and normalizing the cumulative activity to the absolute cellular protein mass, according to the following:

Total Activity (RLU)=(Mean RLU×50)+(Mean RLU×10)

Total Cellular Protein (μg)=Lysate Concentration (μg μL^-1^)×800 μL

The final specific reporter activity was expressed as Total RLU per μg of total protein.

### Microscopy

Image acquisition, with an exposure time of 150 ms, was executed using a Zeiss Axio Imager M1 (Oberkochen, Germany) integrated with the MetaMorph software package (version 7.7.0.0, Molecular Devices, Seattle, IL, USA). Region of interest, addition of scale bars and application of Fire LUT for PI fluorescence images was carried out using (FIJI is just) ImageJ 1.54p, developed by Wayne Rasband and contributors, National Institutes of Health, USA.

### Maize infection assay

Maize (*Zea mays L*.) variety Early Golden Bantam was used for all infections. *U. maydis* strain used for virulence testing included SG200 wildtype, the SG200-TIN2 knockout and optogenetic strains. All strains were cultured in YEPS medium at 28 °C with shaking at 160 rpm. For light-treated cultures, flasks were illuminated with blue light at an intensity of 10 µmol/m²·s unless otherwise specified; dark-treated cultures were wrapped in aluminum foil. When the optical density at 600 nm (OD600) reached 0.8, cells were harvested by centrifugation at 5,000 × *g* for 10 min, washed, and resuspended in sterile distilled water to a final OD600 of 1.0. Infections were performed by injecting the cell suspension into the stems of 7-day-old maize seedlings using a 1-mL syringe with a needle, approximately 1 cm above the soil. Plants were maintained under controlled growth conditions with a 16-h light/8-h dark photoperiod at 28 °C. Disease symptoms were evaluated at 6-12 d post-infection (dpi).

### Luciferase kinetics of the infected leaves

Leaf disks were cut and soak into a 96-well white flat-bottom plates (Costar) for determination of luciferases activity. Each well has 0.2 ml PBS, 0.01%triton-X100 and 5 mM D-luciferin. Three biological replicates of each experiment were done with four technical replicates each. The plate was sealed with an optically clear film (Sarstedt) and recorded the Luminescence in a Berthold Centro XS3 LB960 microplate reader every 30 minutes (0.1 s integration time). The illumination was set by a customized LED on top of the platereader.

### RNA extraction, cDNA synthesis and RT-qPCR

We collected 2-3cm leaf tissue of 1cm below the injection point, flash froze it in liquid nitrogen and ground it using a TissueLyser II (Qiagen). We carried out RNA extraction using RNeasy kits (Qiagen), using buffer RLC and β-mercaptoethanol. To increase the RNA concentration, we reduced the elution volume to 30 μL and used the flow through to elute once again. We treated the collected RNA sample with DNase (Invitrogen) to digest DNA contamination.

We synthesized cDNA using Superscript II (Invitrogen) using 2 μg of RNA per sample. We carried out qPCRs on a 96-well machine (qTOWER³G, Analytik Jena, Germany), with samples loaded in technical triplicates. Each well contained 1 μL of 10× diluted cDNA and 9 μL of PCR mix consisting of 5 μL of SYBR Green I Master Mix (Promega), 1 μL of forward primer (10 μM stock), 1 μL of reverse primer (10 μM stock), and 0.5 μL of nuclease-free water. RT-qPCR primers used in this study are listed in Table S3. We used the following cycling protocol: pre-incubation at 95°C for 5mins, followed by 40 cycles of 20s at 95°C, 20s at 60°C, 20s at 72°C and then melting curve analysis. We calculated ratios using the geometric means of the Ct values for the gene of interest and the reference genes Oppi and zmGAPDH.

### Statistics and reproducibility

Statistical analyses were performed using Explot (https://github.com/beyerh/ExPlot). Quantitative data are presented as mean ± SEM with n = 3–4 biological replicates per condition; figure legends provide further specification as necessary. Unless otherwise stated, biological replicates represent independent treatments in separate wells (in vitro assays) or on leaves from separate plants. All micrographs, gels, and blots are representative images from at least 2 independent repeated experiments. Statistical significance was computed using t-test or ANOVA with Tukey post hoc tests (multiple comparisons correction), as indicated in figure legends. P < 0.05 were considered statistically significant.

## Acknowledgements

We are grateful to Ute Meyer (Köln, Germany) for assisting with *Ustilago* virulence injection, H.M. Beyer (Düsseldorf, Germany) for the designing of LPA illumination device, S. Kuschel and R. Schönle, TAs from the microbiology institute (Düsseldorf, Germany) for valuable experimental assistance.

## Funding

This work was supported by the Deutsche Forschungsgemeinschaft (DFG, German Research Foundation) SFB1535 (project no. 458090666) to M.D.M, M.F. and M.D.Z, and under the Germany Excellence Strategy (CEPLAS-EXC-2048 project no. 390686111) to G.D. and M.D.Z.

## Author contribution

Conceptualization: MDZ, MF, GD

Methodology: KT, MDM, LH, MDZ

Data Acquisition: KT, MDM, LH, WZ, AR, NH, JP, LW

Supervision: MDZ, MF, GD

Writing: KT, MDZ

Funding Acquisition: MF, MDZ

## Notes

### Competing Interest Statement

The authors have declared no competing interest.

