## Supplementary Material for "LUstiGE, Light responsive Ustilago maydis Gene Expression: Optogenetic control of morphogenesis and pathogenesis in the corn fungal pathogen *Ustilago maydis*"

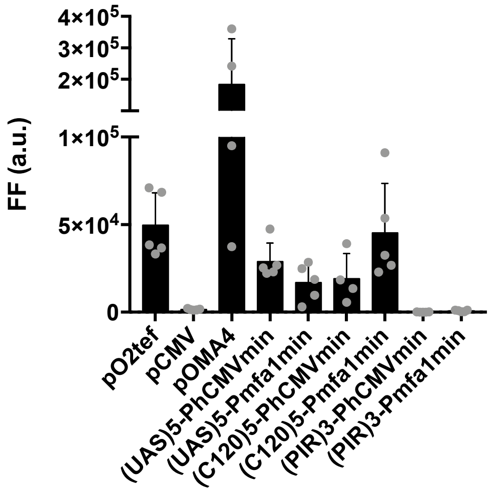


**Figure S1. Comparison of synthetic (orthogonal) promoters for *U. maydis*.** The various synthetic promoters established in this work have been tested for their strength compared to $P_{O2tef}$ (cultures OD600 = 0.5 were analyzed for their FLuc luminescence. Error bars represent the SEM of this individual experiment with n=5).

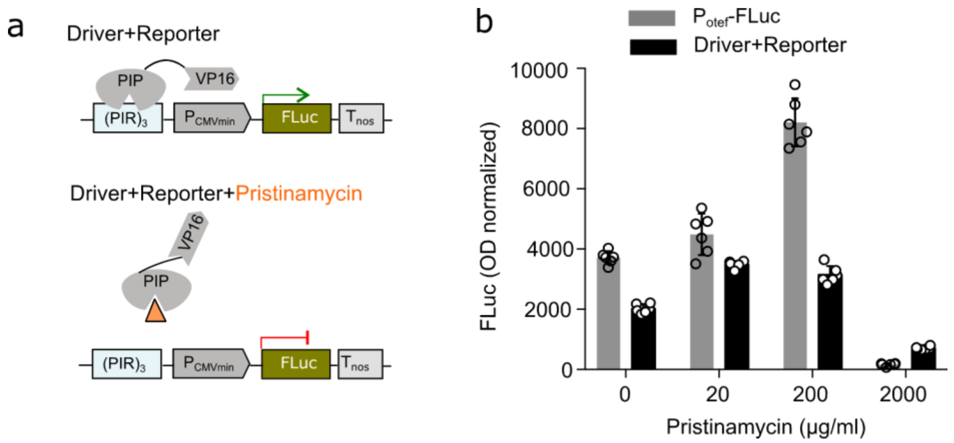


**Figure S2. Dose dependency of PIP/(PIR)3 system. (a)** The PIP-based, synthetic, chemically regulatable gene expression system is encoded on two vectors (Driver+Reporter). The reporter comprises the DNA binding protein PIP fused to the VP16 transactivation domain and an NLS, under the control of a constitutive promoter. The Driver vector carries the PIR3 operating sequence upstream of a minimal promoter, controlling a goi. PIP binds to its operator sequence, which brings the VP16 in close proximity to the minimal promoter, activating the expression of the goi. Upon addition of pristinamycin binding of PIP is inhibited by the antibiotic, and thus expression of the goi stops. Removal of pristinamycin reactivates the expression. (b) cultures of the indicated strains with an OD600 = 0.5 were analyzed for their FLuc luminescence after growing for 24 h on CM-Glucose supplemented with indicated amounts of pristinamycin. The amount of DMSO for a concentration of 2,000 µg ml$^{-1}$ pristinamycin seems toxic as these cultures contained almost no viable cells. Error bars represent the SEM of this individual experiment with n=6.

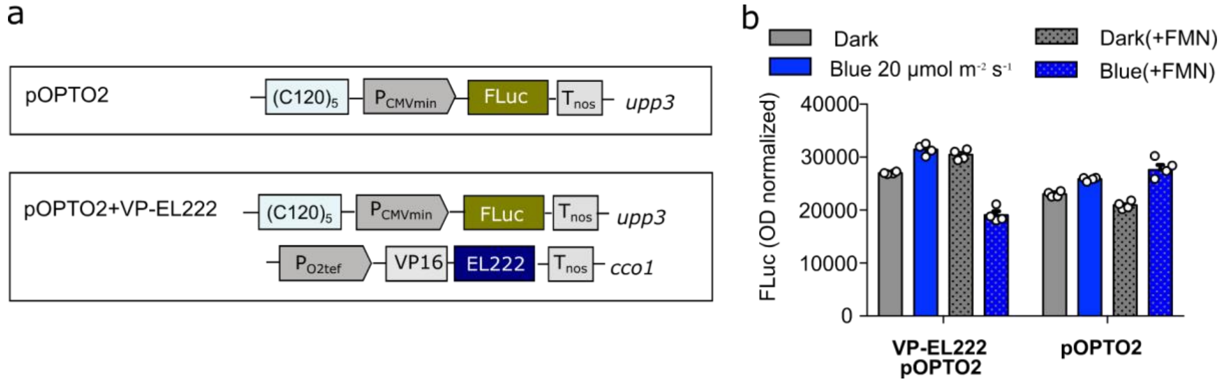


**Figure S3. Functional characterization of the VP-EL222 system in *U. maydis* cultures.** **(a)** Construct design of the EL222 based blue light-controlled repression of gene expression system in *U. maydi*s. Strain pOPTO2+VP-EL222 was generated with two components stably integrated: i) in the *upp3* locus, $(C120)_5$- $P_{CMV}$ minimal promoter, driving the expression of firefly luciferase reporter (FLuc) (pOPTO2); and ii) in the *cco1* locus, the blue light photoreceptor EL222 is fused to a transactivator domain, VP16, and placed under the control of a constitutive $P_{O2tef}$ promoter. A control strain is with only the reporter construct pOPTO2 being integrated to the *upp3* locus. **(b)** Determination of reporter activity of VP-EL222 cultures and control strains were illuminated with blue light for 6 h at 20 µmol $m^{-2}$ $s^{-1}$ light ($\lambda$=450 nm) or kept in the dark. Error bars represent the SEM of this individual experiment with n=4.

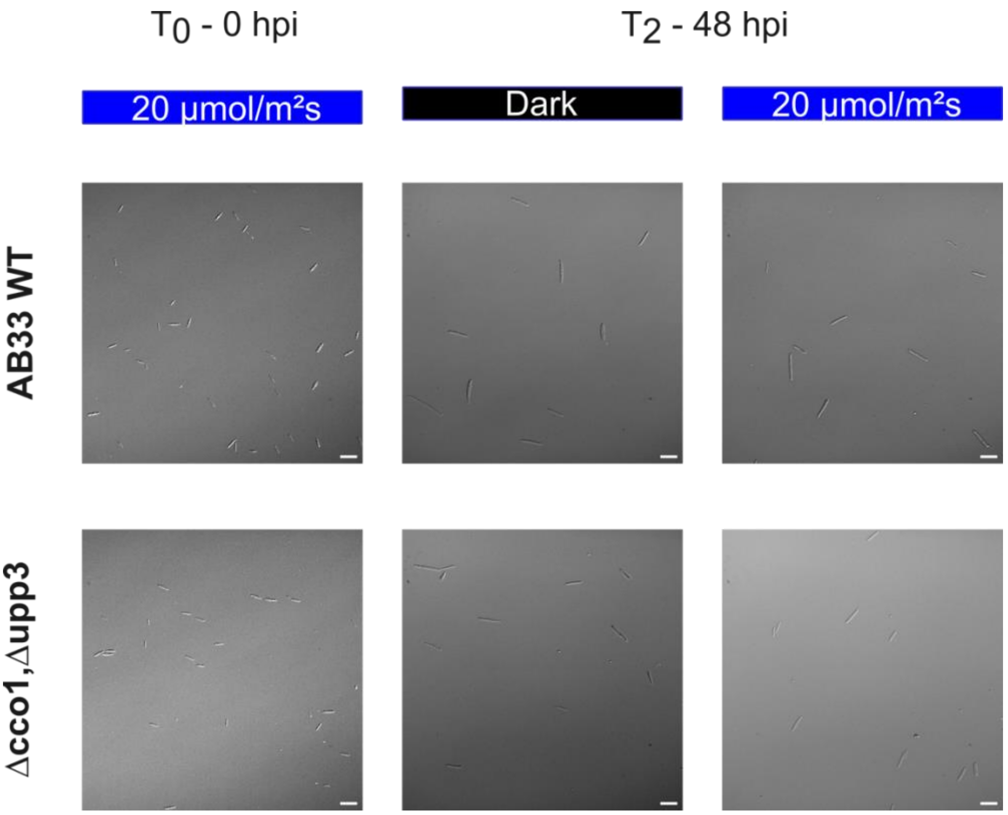


**Figure S4. Optogenetic control of Rac1$^{Q61L}$ expression**
Displayed are DIC images of the negative control strains of figure 4: the wild-type AB33 strain and the genetic background strain (sUma2549) of the Blue-OFF system with its deletions at the Δcco1 and Δupp3 loci. Control samples were evaluated alongside the experimental Rac$^{Q61L}$ culture. Samples from blue-light- (20 µmol m$^{-2}$ s$^{-1}$) and dark-treated control cultures were taken at 0 h ($T_0$), 24 h ($T_1$), 48 h, ($T_2$), and 72 h ($T_3$) (for comparison with the re-illuminated dark-treated cultures of Rac1$^{Q61L}$). At all given time points cell populations were observed as sporidia. Scale bar = 20 µm.

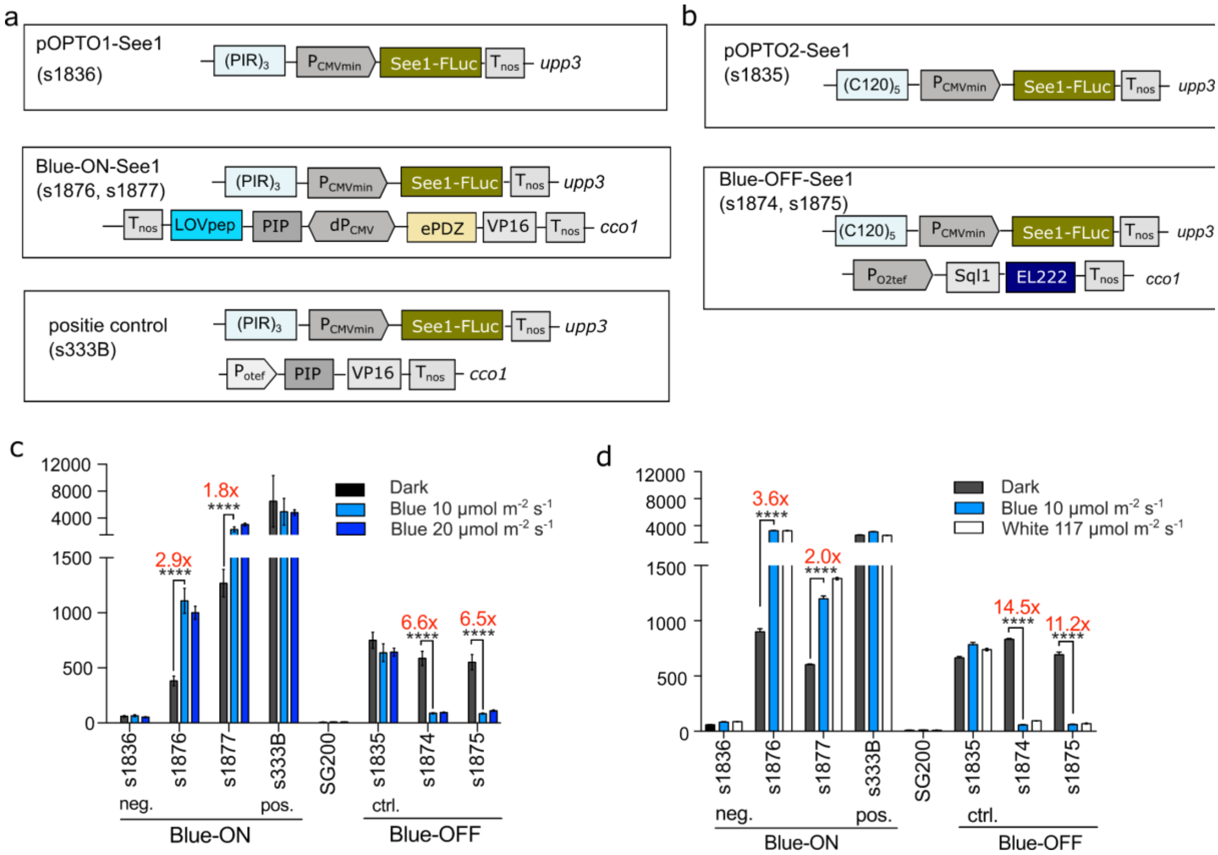


**Figure S5. Optogenetically control of See1 expression.** (A) Construct design for Blue-ON-See1, and (B) Blue-OFF-See1 strains. (C) *U. maydis* cultures were incubated under different light conditions. (C) White light illumination induced the same expression as blue light. Error bars represent the SEM. Statistical significance was calculated using the Student's *t*-test. n.s., not significant, $^{**}p < 0.01$ and $^{***}p < 0.001$.

**Table S1. Generation and description of plasmids used in this work.**

All plasmids are constructed with T4 ligation or AQUA cloning[55] if not indicated otherwise. Grey and bold: plasmids used in strain generation; white: intermediate cloning plasmids.

| **Plasmid** | **Description** |
|---|---|
| pLH112 | **(tetO)13-$P_{CMVmin}$-FLuc-Tnos** |
| pLH113 | **(tetO)13-$P_{mfa1min}$-FLuc-Tnos** |
| pLH120 | **(C120)5-$P_{CMVmin}$-FLuc-Tnos** |
| pLH121 | **(C120)5-$P_{mfa1min}$-FLuc-Tnos** |
| pNH019 | **(UAS)5-$P_{CMVmin}$-FLuc-nosT** |
| pNH021 | **(UAS)5-$P_{mfa1min}$-FLuc-Tnos** |
| pNH023 | **(PIR)3-$P_{CMVmin}$-FLuc -Tnos** |
| pNH025 | **(PIR)3-$P_{mfa1min}$-FLuc -Tnos** |
| pLH126 | **PO2tef-NLS-Sql1-EL222-Tnos** |
| pLH107 | **PO2tef-TetR-VP16ff-NLS-Tnos** |
| pLH111 | **PO2tef-NLS-VP16ff-EL222-Tnos** |
| pNH006 | **PO2tef-GAL4BD-VP16ff-NLS-Tnos** |
| pNH008 | **PO2tef-PIP-VP16ff-NLS-Tnos** |
| pNH047 | **Tnos-ePDZ-VP16ff-NLS::dPhCMV::PIP-LOVpep-Tnos** |
| pMDM13 | **(C120)5-$P_{CMVmin}$-Rac1$^{Q61L}$-Tnos** |
| pOPTO1-See1 | **PIR3-$P_{CVMmin}$-See1-Fluc-Tnos** |
| pOPTO2-See1 | **(C120)5-$P_{CMVmin}$-see1-FLuc-Tnos** |
| pKT1421 | **PIR3-$P_{CVMmin}$-TIN2-Tnos** |

**Table S2. Strains used in this work.**

| Strains | Description / Genetic background | **Transformed Plasmid*** | **Origin** |
|---|---|---|---|
| sLHNH008 | AB33_upp3D::$P_{O2tef}$::FLuc-HA-Tnos-NatR | | Heucken et al.,2023[2] (Fig. S1) |
| | AB33_upp3D::$P_{mfa1}$::FLuc-Tnos-NatR | | This work (Fig. S1) |
| sNH026 | AB33_upp3D::$(UAS)_5$-$P_{CMVmin}$::FLuc-Tnos-NatR | pNH019 | This work (Fig. S1) |
| sNH028 | AB33_upp3D::$(UAS)_5$-$P_{mfa1min}$::FLuc-Tnos-NatR | pNH021 | This work (Fig. S1) |
| sNH030<br>pOPTO1 | AB33_upp3D::$(PIR)_3$-$P_{CMVmin}$::FLuc-Tnos-NatR | pNH023 | This work (Fig. S1, Fig.2) |
| sNH032 | AB33_upp3D::$(PIR)_3$-$P_{mfa1min}$::FLuc-Tnos-NatR | pNH025 | This work (Fig. S1) |
| sLH008 | AB33_upp3D::$(tetO)_{13}$-$P_{CMVmin}$::FLuc-Tnos-NatR | pLH112 | This work (Fig. S1) |
| sLH009 | AB33_upp3D::$(tetO)_{13}$-$P_{mfa1min}$::FLuc-Tnos-NatR | pLH113 | This work (Fig. S1) |
| sLH023<br>Blue-ON | AB33_cco1D::Tnos-ePDZ-VP16ff-NLS::dPhCMV::PIP-LOVpep-Tnos-HygR_upp3D::$(PIR)_3$-$P_{CMVmin}$:: FLuc-Tnos-NatR | pNH047<br>pNH023 | This work (Fig. 2) |
| sNH056<br>positive control | AB33_cco1D:: PO2tef::PIP-VP16ff-NLS-Tnos-HygR_upp3D:: $(PIR)_3$-$P_{CMVmin}$::FLuc-Tnos-NatR | pNH008<br>pNH023 | This work (Fig. 2) |
| sLH016<br>pOPTO2 | AB33_upp3D::$(C120)_5$-$P_{CMVmin}$::FLuc-Tnos-NatR | pLH120 | This work (Fig. 3) |
| sLH017 | AB33_upp3D::$(C120)_5$-$P_{mfa1min}$::FLuc-Tnos-NatR | pLH121 | This work (Fig. 3) |
| sLH020 | AB33_cco1D::$P_{O2tef}$::VP16ff-EL222-Tnos-HygR_upp3D::C120-$P_{CMVmin}$:: FLuc-Tnos-NatR | pLH111<br>pLH120 | This work (Fig. 3) |
| sLH026 | AB33_cco1D::PO2tef::NLS-Sql1-EL222-Tnos-HygR_upp3D::$(C120)_5$-$P_{CMVmin}$:: FLuc-Tnos-NatR | pLH126<br>pLH120 | This work (Fig. 3) |
| sMDM36.8<br>Blue-OFF-Rac1$^{Q61L}$ | AB33_cco1D::PO2tef::NLS-Sql1-EL222-Tnos-HygR_upp3D::$(C120)_5$-$P_{CMVmin}$::Rac1$^{Q61L}$-Tnos-NatR | pLH126<br>pMDM13 | This work (Fig. 4) |
| SG200 | | | This work (Fig. 5,6) |
| **s1835**<br>pOPTO2-See1 | SG200_upp3D::$(C120)_5$-$P_{CMVmin}$::See1-FLuc-Tnos-NatR | pOPTO2-See1 | This work (Fig. S3, Fig.5) |
| **s1836** | SG200_upp3D::$(PIR)_3$-$P_{CMVmin}$::See1-FLuc-Tnos-NatR | pOPTO1-See1 | This work (Fig. S3,) |
| **s1874, s1875**<br>**Blue-OFF-See1** | SG200_cco1D::$P_{O2tef}$::NLS-Sql1-EL222-Tnos-HygR _upp3D::$(C120)_5$-$P_{CMVmin}$::See1-FLuc-Tnos-NatR | pLH126<br>pOPTO2-See1 | This work (Fig. S3, Fig.5) |
| **S1876, s1877**<br>**Blue-ON-See1** | SG200_cco1D::Tnos-ePDZ-VP16ff-NLS::dPhCMV::PIP-LOVpep-nosT-HygR_upp3D::$(PIR)_3$-$P_{CMVmin}$::See1-FLuc-Tnos-NatR | pNH047<br>pOPTO1-See1 | This work (Fig. S3) |
| **s333B**<br>**positive control** | SG200_cco1D:: $P_{O2tef}$::PIP-VP16ff-NLS-Tnos-HygR_upp3D::(PIR)3-$P_{CMVmin}$::See1-FLuc-Tnos-NatR | pNH008<br>pOPTO1-See1 | This work (Fig. S3) |

| | | | |
|---|---|---|---|
| **s798** | SG200-ΔTIN2 | | This work (Fig. 6) |
| **s2575** | SG200-ΔTIN2_cco1D::Tnos-ePDZ-VP16ff-NLS::dPhCMV::PIP-LOVpep-Tnos-HygR | pNH047 | This work (Fig. 6) |
| **s2583**<br>**Blue-ON-TIN2** | SG200-ΔTIN2_cco1D::Tnos-ePDZ-VP16ff-NLS::dPhCMV::PIP-LOVpep-Tnos-HygR_upp3D::PIR3-$P_{CMVmin}$::TIN2-Tnos-NatR | pNH047<br>pKT1421 | This work (Fig. 6) |

### Table S3. oligonucleotides used for RT-qPCR in this work.

| **primer** | | Sequence(5-3) |
|---|---|---|
| **oROF424** | **Fluc** | GAGGCGAACTGTGTGTGAGA |
| **oROF425** | | GTGTTCGTCTTCGTCCCAGT |
| **oKT398** | **Oppi** | ACATCGTCAAGGCTATCG |
| **oKT399** | | AAAGAACACCGGACTTGG |
| **oKT396** | **See1** | TCAGGTGCAAGGAGAAGG |
| **oKT397** | | ACAGAATACTCCGCTTCCC |
| **oKT795** | **TIN2** | TGCCATCATTTCGACCCCAA |
| **oKT796** | | GAAGCGAGGGAGCAGATTGT |
| **oKT789** | | GATGTACCGCCGCAAGATGG |
| **oKT790** | **ZmFHT1** | GCAAGAATGGCGTCGAGAGG |
| **oKT791** | | TGGACCTGGTCACCATCATC |
| **oKT792** | **ZmDFR** | ATGAGCTGCACCTGCTTGAG |
| **oKT789** | | GATGTACCGCCGCAAGATGG |
| **oKT790** | **ZmFHT1** | GCAAGAATGGCGTCGAGAGG |
| **oKT394** | | GAATCAACGGCTTCGGAAGGAT |
| **oKT395** | **ZmGAPDH** | CCTCAGGGTTCCTGATGCCAAA |
